# Dissolved oxygen dynamics reveal biogeochemical tipping points driven by river corridor hydrology

**DOI:** 10.1101/2020.01.31.929505

**Authors:** Matthew H Kaufman, Ruby N Ghosh, Jay Grate, Dean D Shooltz, Michael J Freeman, Terry M Ball, Reza Loloee, Charles W McIntire, Jackie Wells, Chris Strickland, Vince Vermeul, Kenton A. Rod, Rob Mackley, Xinming Lin, Huiying Ren, James Stegen

**Author notes:** Matthew H Kaufman and Ruby N Ghosh contributed equally to this manuscript.

## Abstract

Riverbeds can have an important impact on large-scale fluxes of biogeochemically active solutes in river corridor systems. The hyporheic zone is an important area of both hydrology and biogeochemistry, particularly in a hydropeaked river system where rapid variations in river stage height, hydraulic gradients, and residence times occur. We measured several biogeochemical and hydrological parameters at three different subsurface depths in the riverbed of the Columbia River in Washington state, a complex hydropeaked river system with significant subsurface heterogeneity. During the study, episodes of significant dissolved oxygen (DO) change were observed. The DO signal changes were the most apparent, compared to more modest changes in other biogeochemical markers. While DO is often associated with biological activity, we ultimately found that the notable DO excursions were associated with hydrologic gradients. Here we describe hydrologic perturbations and biogeochemical responses in terms of hydrobiogeochemical regimes. Two different forms of DO response to hydraulic gradient perturbation were observed, defining different hydrobiogeochemical regimes. The system tips abruptly from one regime to the other, exhibiting threshold behavior that is uncaptured in current estimations of cumulative influences of subsurface processes from reach to earth system scales.

**Plain language summary:** The water in a river is constantly flowing in and out of the river’s bed. While it is in the bed, microbes process environmentally and ecologically important nutrients and other compounds. The exchange of water between the river and the riverbed is largely controlled by the stage of the river. Many processes, both natural and human, can cause rapid changes in the river stage. These rapid stage changes can cause abrupt changes in the riverbed’s ability to process nutrients and other compounds. In this study, we use dissolved oxygen as a window into riverbed microbial and chemical processing. We see that relatively rapid changes in river stage can cause abrupt changes in the concentrations of chemicals in the riverbed, indicating fundamental changes in how the riverbed processes are occurring. Understanding these abrupt and fundamental changes is important to our overall understanding of the ecological function of river corridor systems.

**3 key points:**

1. A hydrobiogeochemical regime is defined by the functional form of the relationship of a hydrologic perturbation and biogeochemical marker.
2. Short-term hydrologic perturbations drive tipping points between distinct hydrobiogeochemical regimes.
3. Linear and hysteretic dissolved oxygen dynamics define distinct regimes as observed throughout the hyporheic zone.

## 1 Introduction

Riverbeds play host to important biogeochemical processes [*Boano, et al*., 2015, *Boulton, et al*., 1998, *Brunke and Gonser*, 1997, *Fischer, et al*., 2005]. The biogeochemical conditions present at any given time and location in the riverbed are a function of both reaction rates and residence times [*Zarnetske, et al*., 2011, *Zarnetske, et al*., 2012]. Rivers are also highly dynamic, driving subsequent dynamics in reaction rates [*Mcclain, et al*., 2003] as well as flowpaths and residence time distributions [*Gomez-Velez, et al*., 2017, *Gomez and Wilson*, 2013, *Larsen, et al*., 2014]. Hydrologic dynamism in rivers occurs over a wide range of timescales [*Boano, et al*., 2015, *Kirchner and Neal*, 2013, *Woessner*, 2000]. Increasing attention has been paid to the impact of short term hydraulic perturbations, on the hours-to-days scale, such as those created by hydropeaking, tidal forces, evapotranspiration, and other highly dynamic processes [*Singh, et al*., 2019, *Wolke, et al*., 2020]. These intricately linked and highly dynamic processes combine to control the concentrations of chemical markers related to biogeochemistry in the riverbed, particularly electron donors and acceptors [*Stegen, et al*., 2018].

In this paper we report the results of monitoring dissolved oxygen (DO) at subsurface locations in the hyporheic zone at a field site along the Columbia River. The optical DO sensor was deployed in a shoreside location with sensors for several other chemical and physical parameters, in a measurement system that drew samples from a dense array of aquifer tubes. In this study, water was sampled from three locations at different depths in the subsurface riverbed. During the period of observations (16 days), episodes of significant DO change were observed as “apparent hot moments”. The DO signal changes were the most apparent, compared to more modest changes in other parameters such as nitrate, ORP, and salinity; without these DO excursions, nothing of significance may have been noted during this time period. While DO is a biogeochemical parameter often associated with biological activity, we ultimately found, from pressure readings, that the notable DO excursions were associated with hydrologic gradients. Hydrologic gradients and flowpaths, in turn, are expected to influence the biogeochemistry. These events occurred as part of a complex hydropeaked river system with significant subsurface heterogeneity.

Ultimately, perturbations of biogeochemical markers like oxygen are driven by varying ratios of reaction rate timescale to transport timescale, a ratio described by the Damkohler number [*Damköhler*, 1936, *Zarnetske, et al*., 2012]. Traditionally, such perturbations have been named “hot moments” if they meet the criteria laid out by *Mcclain, et al*. [2003]: “biogeochemical … hot moments are defined as short periods of time that exhibit disproportionately high reaction rates relative to longer intervening time periods”. However, in highly dynamic and spatially heterogeneous systems, variations in flowpaths and residence times can lead to similar perturbations of biogeochemical markers, independent of changes in reaction rate [*Briggs, et al*., 2014]. In order to avoid confusion, here we refer to a particular functional form of hydrologic perturbation and biogeochemical response as a “hydrobiogeochemical regime”. This terminology provides the ability to examine the various ways that biogeochemical markers respond to hydraulic perturbations without being ambiguous or limited to specific mechanisms, as is the case with a strictly defined hot moment. We can also apply the term more broadly. Where in this particular scenario, the hydrobiogeochemical regime is defined by the functional form of the relationship between a biogeochemically active solute and hydrologic gradient, the details of those axes can be different across studies. For example, the functional form of a relationship between nitrate and vertical gradient could also define a hydrobiogeochemical regime. Ultimately, this implies that there are many hydrobiogeochemical regimes that are co-existing in time. This is conceptually similar to metabolic regime, a term defined by the relationship between ecosystem respiration and gross primary productivity [*Bernhardt, et al*., 2018]. Both relate two dynamic variables that are causally linked to each other and use the functional relationship to define some property of the regime.

Biogeochemical responses to hydrologic perturbations are often attributed to variable mixing of oxic surface water and suboxic groundwater [*Arntzen, et al*., 2006, *Mcallister, et al*., 2015], particularly through the mechanism of variable reaction rates driven by changing microbial access to nutrients, organic matter, and electron acceptors as those mixing ratios change [*Marzadri, et al*., 2016]. Outside of mixing however, variable stage elevations can lead to rapid changes in lateral hydraulic gradients [*Fritz and Arntzen*, 2007, *Watson, et al*., 2018], which have the potential to drastically alter residence time distributions and access to preferential flowpaths [*Poole, et al*., 2006, *Sawyer, et al*., 2009]. While several studies conclude that changes in residence time and flowpath length drive biogeochemical responses [*Briggs, et al*., 2014, *Valett, et al*., 1996], there is currently relatively little research into biogeochemical responses to short term hydraulic perturbations in field locations with low levels of groundwater/surface water mixing. This results in a limited ability to isolate gradient-driven, residence-time based effects from mixing-driven, reaction-rate based effects.

The variable inundation and variable lateral gradient created by a river with rapid stage fluctuation, combined with heterogeneity in subsurface hydrologic properties presents the potential for strongly non-linear biogeochemical responses, and offers the possibility for rapid changes between hydrobiogeochemical regimes as preferential flowpaths and subsurface hydraulic connectivity vary in response. In this study, we hypothesize that in the absence of strong groundwater/surface water mixing, dissolved oxygen dynamics will be explained primarily by variations in hydraulic gradients. Additionally, we hypothesize that stage dynamics will lead to potentially abrupt changes between hydrobiogeochemical regimes. We observe hydrobiogeochemical dynamics during a period of little groundwater/surface water mixing, and are able to explain a large portion of the dissolved oxygen variation using only hydraulic gradients. This implies that variations in residence time can drive biogeochemical dynamics independently of mixing. Additionally, we observe abrupt changes in the functional relationship between hydrologic drivers and biogeochemical responses. These threshold-like changes in hydrobiogeochemical regime in the subsurface are likely a result of gradient and stage driven flowpath activation/deactivation, influenced by subsurface heterogeneity.

## 2 Methods

### 2.1 Field monitoring system

The field setting for this study was the shore and near-shore environment of the Columbia River at the 300 area of the Hanford site near Richland, Washington (Figure 1). The Hanford reach of the Columbia River is contained within the Pasco Basin in southeast Washington State. The river is surrounded by an unconfined aquifer consisting of fluvial Ringold Formation and Hanford Formation flood sediments [*Hartman and Dresel*, 1998]. The permeability of the riverbed in this area is highly variable, ranging from 2.8×10^−5^ cm/s to 4.3×10^−2^ cm/s [*Arntzen*, 2001].

**Figure 1:**
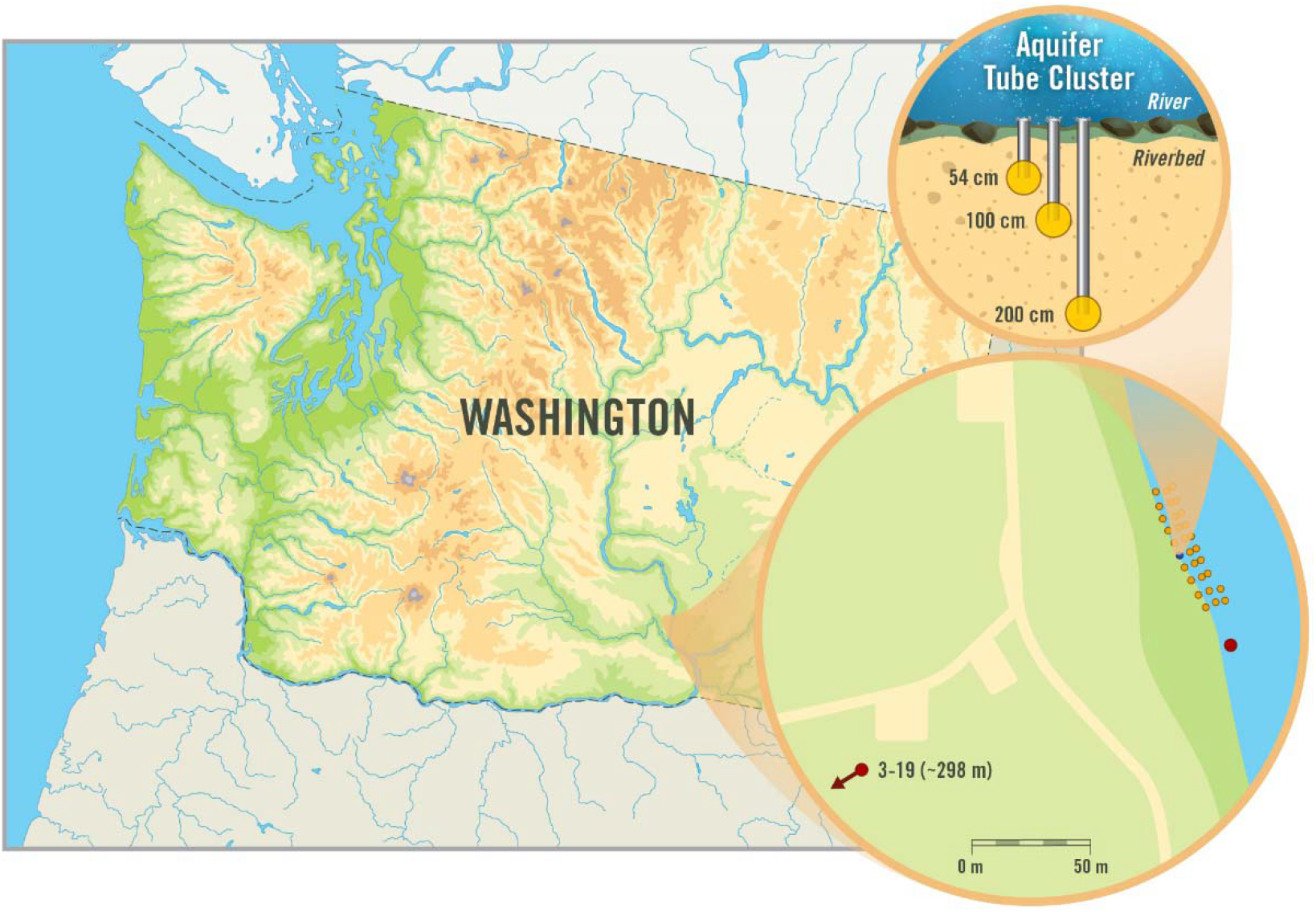
Field site location near Richland WA, USA. Dense array aquifer tube clusters are indicated with orange dots, other wells and piezometers are indicated with red dots.

Subsurface biogeochemistry was monitored from April 7^th^ to April 23^rd^ 2018 using a small subset of the aquifer tubes installed in the “dense array”, a unique hyporheic zone water sampling instrument which takes real time water chemistry measurements from a large array of riverbed monitoring points. (Figure 1). This instrument included an array of linear low density polyethylene tubes (12.7 mm [½″] outer diameter, 6.35 mm [¼″] inner diameter) installed in a grid pattern along the permanently wetted riverbed along the shoreline. The tubes were installed in clusters and were spaced over an approximately 10 m (perpendicular to the shoreline) by 90 m (parallel to the shore) grid 10 clusters long and three clusters wide. Each cluster consisted of 2 or 3 tubes installed to approximate depths of 0.5 m, 1 m, 2m below the sediment-water interface, allowing for three dimensional sampling of the hyporheic zone. Due to substrate heterogeneities some locations could not be penetrated with the hand installation tools used and as a result not all clusters contain a 2+ meter monitoring point. The end of each tube was screened to form a mini-piezometer, with slots cut into the last 3 inches to 6 inches of each tube to allow water to be sampled. These slots were covered with Teflon mesh to reduce sediment clogging. The ends of the tubes were closed with stainless steel thermistor (Figure 2). Upon installation, each tube was developed by placing each tube under 0.6-0.8 atmosphers of vacuum until 500 ml of water had bee withrdrawn, then pumping water into the tube for 1 minute, and again withdrawing 500ml of water under vacuum. This process was repeated until clarity of the water withdrawn under vacuum stabilized.

**Figure 2:**
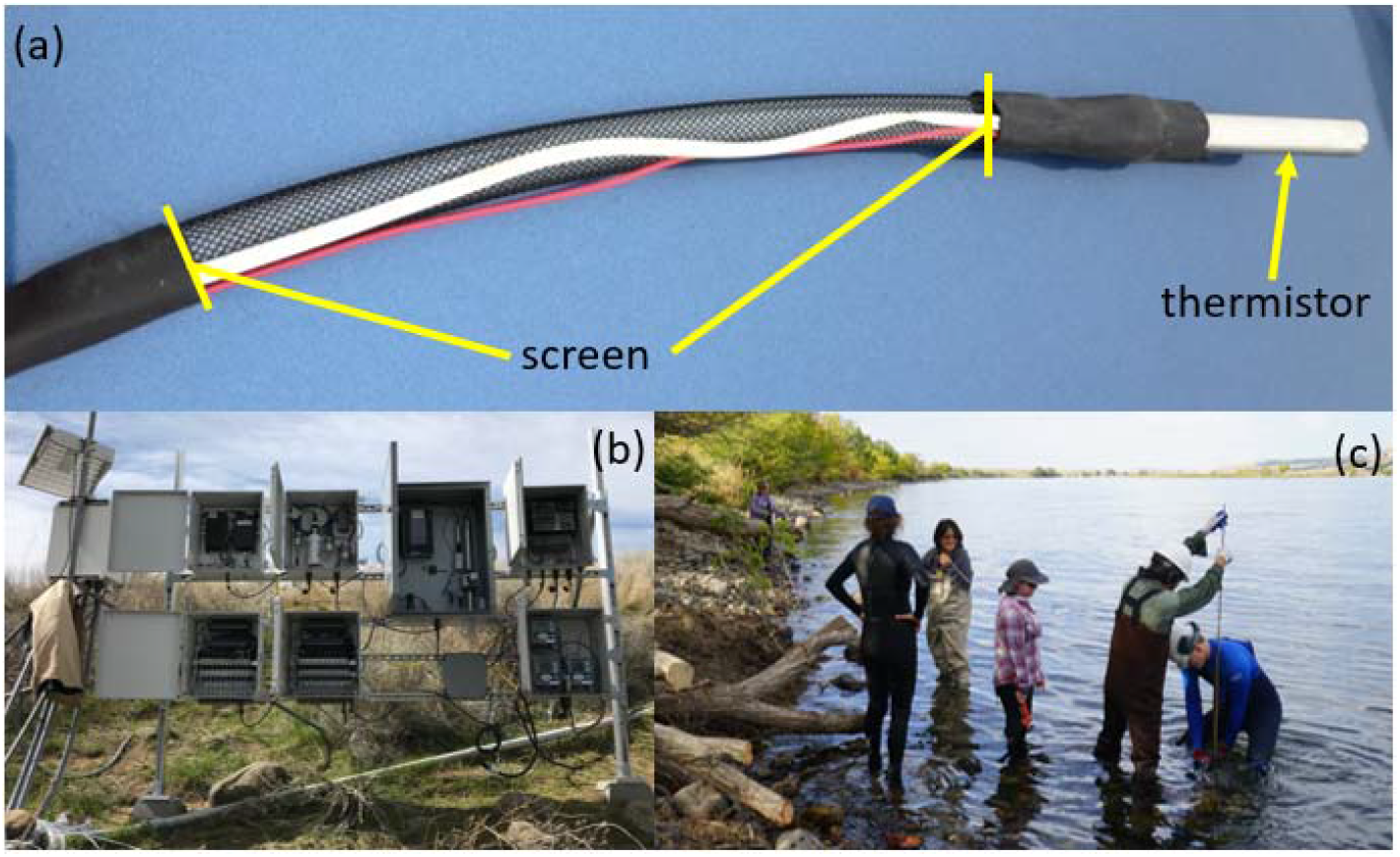
(a) shows the tip of the aquifer tubes, with the screened section and thermistor cap. (b) shows the series of manifolds and datalogging system. (c) shows the hand-installation of an aquifer tube.

Each of the sampling tubes was connected to its own solenoid valve, with all valves being controlled by a CR3000 datalogger through four SDM-CD16AC 16-Channel AC/DC Relay Controllers (Campbell Scientific, Logan UT). In typical operation, the solenoid valves were opened one at a time for 20 minutes each and two coupled Kloehn syringe pumps (IMI Precision Engineering, Las Vegas NV) drew sample water from the tube and delivered it to the monitoring manifold at a rate of 5 mL s^−1^.

Our focus here is to obtain high time resolution sampling from multiple depths at a single x-y location in the dense array (Figure 1). By focusing on one vertical transect, the 40min pumping time per solenoid allowed each depth to be sampled every two hours. The pumps delivered sample water to a series of manifolds containing several instruments, for real time measurements of dissolved oxygen and water chemistry parameters (Figure 2).

The time resolved dissolved oxygen measurements were acquired with a novel optical DO sensor developed by OptiO_2_, LLC (www.optio2.com), incorporated into the dense array system with a flow cell housing the DO probe (Figure 3a). DO was recorded every 30 seconds as the solenoid valves cycled between sampling tubes collecting stream water from depths of 54cm, 100cm, and 200cm (Figure 3b). This high temporal resolution was necessary to correlate measured DO with pore water sampling depth. Our experimental conditions (40 minutes of pumping at each depth with DO recorded every 30 seconds) were sufficient to cleanly separate the data from each location. The OptiO2 probe is a selfcontained optical spectrometer which outputs a temperature and pressure corrected DO signal. Molecular oxygen is detected by monitoring the phosphorescence emission from molybdenum chloride optical indicators [*Ghosh, et al*., 2011]. Ultra-violet photons pump the optical indicators to a spin triplet excited state, which is specifically quenched by the ground state of molecular oxygen, ^3^O_2_. The resultant optical sensing film, which is in direct contact with the liquid to be analyzed, has minimal cross sensitivity to organic and inorganic species and does not suffer from photobleaching [*Ghosh, et al*., 2019]. These properties enabled DO measurements with high sampling rates for an extended period of time from the unconditioned field water samples, with highly variable temperatures and complex chemical constituents.

**Figure 3:**
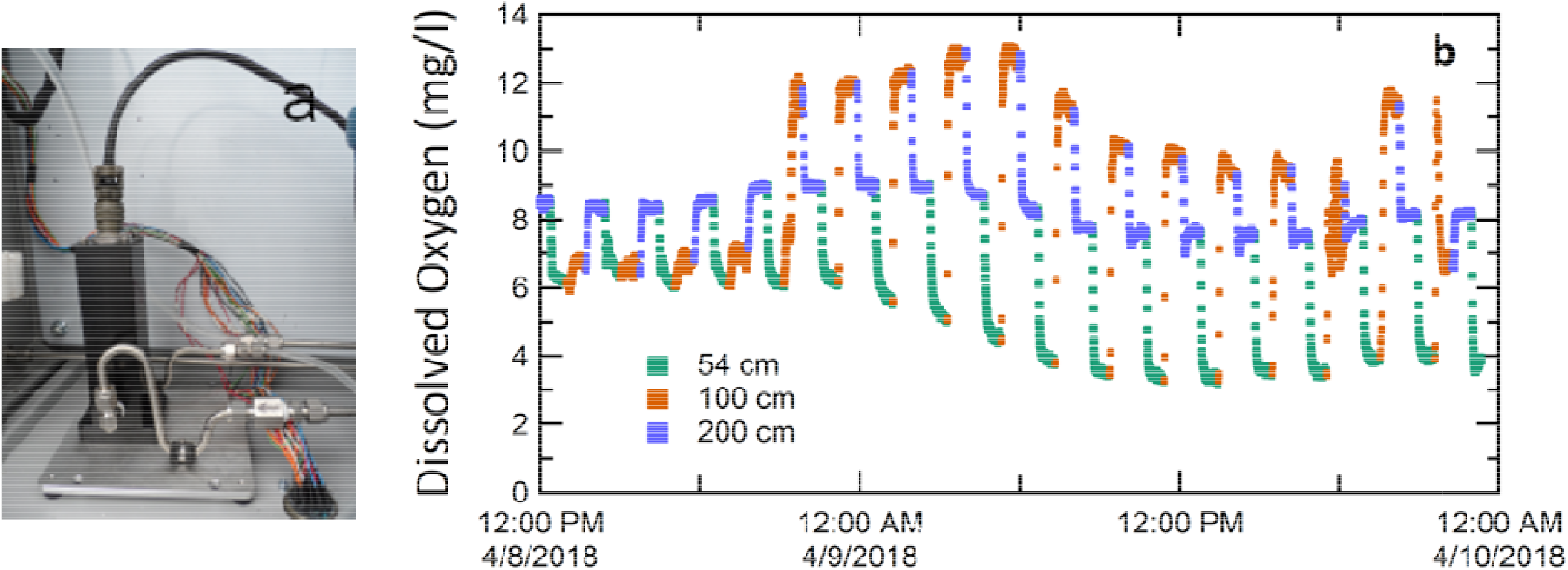
(a) is the Opti O_2_ DO sensor housed in a flow cell incorporated into the dense array manifold. (b) is a 48-hour subset of the raw 30-second DO data recorded by the sensor, showing the variation in DO signal as the dense array switches between the three sampling depths.

In addition to the DO instrument, the pumps delivered sample water to a series of manifolds containing several instruments, for real time measurements of water chemistry (Figure 2). Measurements were made for nitrates using a field S::CAN spectro::lyser UV-Vis 35mm (S::CAN Messtechnik GmbH, Vienna, Austria), pH and redox potential using a S::CAN ph::ilizer/redo::ilizer V2 (S::CAN Messtechnik GmbH, Vienna, Austria), conductivity with Rosemount™ 400 (Emerson, Irvine, CA), pH and oxidation-reduction potential (ORP) with Rosemount™ 3900 (Emerson, Irvine, CA). All data from these instruments were controlled and data was logged by the CR3000. Most instruments were cleaned and maintained biweekly throughout the field season, while the OptiO2 device did not require any recalibration or maintenance.

Immediately downstream of the dense array a transect of five, 3.81 cm (1.5”) inner diameter, stainless steel piezometers were installed. One of the piezometers was left open to the river with the screen at the river level, the other four were capped and had a sensor installed at the screen to allow for depth discreet measurements of hyporheic zone hydraulic head. Of the four capped piezometers, two were installed in the permanently inundated channel with one shallow (P1S) screened at 104.241masl and the other deeper (P1D) screened at 101.67masl. The other two were located on the ephemerally inundated cut bank with one shallow (P2S) screened at 105.689masl and the other deeper P2D screened at 103.884masl. The piezometers were each outfitted with an Aqua TROLL sonde (In-Situ, Inc. Ft. Collins CO) for continuous measurement of pressure, temperature, and conductivity. The sonde measuring the river was vented and the other four sondes were not vented but were corrected for barometric changes in pressure at the data logger. The sondes were connected to a CR1000 data logger (Campbell Scientific, Logan UT) and data was collected twice a month during the field season.

A 10-year (2008--2018) hourly spatio-temporal dataset was collected from a network of groundwater wells at the 300 Area of the U.S. Department of Energy Hanford site, located in southeastern Washington State. Three measurements are monitored at ~40 wells which contains temperature, water elevation and specific conductance. The well network was built to monitor the attenuation of legacy contaminants. The water elevation dynamics in each groundwater well are driven by river stage fluctuations, which in turn influence contaminant recharge to groundwater and lead to highly complex transport behaviors of the contaminants at the site [*Arntzen, et al*., 2006, *Ren, et al*., 2019, *Zachara, et al*., 2016].

### 2.2 Data processing

Data was recorded at 30 second intervals for the analytes in the dense array, and 15 minute intervals for the wells, piezometers, and river channel data. Solenoids switched from one of the three depths to the next every 41 minutes. Because the tubes are relatively long, it takes many minutes for the sampling manifold to come to equilibrium with the incoming sample water after each solenoid switch. To deal with this, the first 31 minutes of each 41 minute monitoring event was discarded, and the data for the final 10 minutes was averaged. This provides one measurement for each analyte for each depth approximately every 2 hours for the duration of the study.

### 2.3 Data analysis

Principal Component Analysis (PCA) is a multivariate statistical method that transforms a large set of possibly correlated initial variables into a smaller number of uncorrelated new variables, called principal components (PCs), which contain most of the variation in the large set [*Ringnér*, 2008]. PCA is mostly used as a tool for exploratory data analysis, dimension reduction, and for making predictive models. It can also be used to evaluate the genetic distance and relatedness between original variables. In this study, we will use PCA to assess the distance and relatedness between gradient variables in order to choose a representative gradient for further analysis.

Mathematically, the PCs are linear combinations of the initial variables weighted by their contribution to explaining the variance in a particular orthogonal dimension. The *i*-th PC is can be calculated as [*James, et al*., 2013]

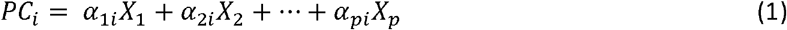

Here *X*_1_, *X*_2_,… *X_p_* are the standardized initial variables, meaning *X_i_* has been centered to have mean zero and standard deviation one. *α*_1*i*_, *α*_2*i*_, … *α_pi_* are the loadings of the *PC_i_*, and 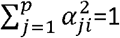.

These PCs reflect both common and unique variance of the variables and may be seen as a variance-focused approach seeking to reproduce both the total variable variance with all components and to reproduce the correlations. The measurement of the variance along each PC also provides a means for comparing the relative importance of each dimension [*Wold, et al*., 1987]. For example, the first PC, which has the large variance, is the most important one among all PCs.

Spearman’s rank correlation is a non-parametric test that used to assess the monotonic association between two ranked variables [*Myers, et al*., 2013]. Unlike Pearson correlation, which measures the linear relationship between normally distributed variables [*Benesty, et al*., 2009], Spearman’s rank correlation test does not carry any assumptions about the distribution of the data. The measure of Spearman’s rank correlation has a value between +1 and −1, where 1 means the ranks of two variables are perfectly monotonically and positively related, −1 means a perfectly monotonic inverse relationship between the two ranked variables, 0 means no correlation between the ranks. When all n ranks are distinct integers, the Spearman rank correlation between two variables can be formulated as [*Zar*, 2005]:

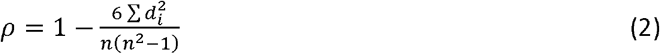

Where ρ is the Spearman’s rank correlation, *d*_1_ is the difference between the ranks of the corresponding variables, and *n* is the number of observations. Otherwise,

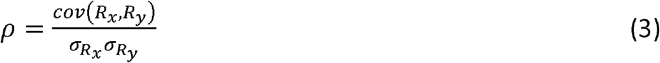

where *cov*(*R_x_*, *R_y_*) is the covariance of the rank variables *R_x_* and *R_y_*, *σ_R_x__* and *σ_R_y__* are the standard deviations of the rank variables [*Zar*, 2005]. In our work, the Spearman’s rank correlation is specifically used to assess the association between biogeochemical markers like dissolved oxygen and nitrate and hydrologic indicators such as hydraulic gradient, specific conductivity, and temperature. A Spearman’s rank correlation matrix [*Chatfield*, 2018] is created to summarized these relationships.

Hydraulic gradients were computed between 4 monitoring wells to the west of the river and the various piezometers located in or near the river. These gradients were approximately horizontal. One vertical gradient was also calculated using the P1S and P1D piezometers. This vertical gradient was highly correlated with the horizontal gradients. Using the entire data set, the first principal component contained only gradients and stage (Table S1). This in part indicates that these hydraulic variables may be highly covariant. Correlation between hydraulic variables was assessed using Spearman rank. All gradients were closely correlated, indicating a consistent regional hydraulic gradient in this area. The gradient computed between background well 3-19 and in-river piezometer P1D was most highly correlated with all other computed gradients (Table 1), and thus was selected as the representative hydraulic gradient in order to simplify all further analyses.

**Table 1:** Correlation between hydraulic gradient at well pair P1D,3-19 and other local well pairs.

| well pair | Spearman Rank | p value |
| --- | --- | --- |
| P1S,P2D | 0.91 | <0.001 |
| P1D,1-2 | 1 | <0.001 |
| P1D,2-3 | 0.98 | <0.001 |
| P1D,2-2 | 0.99 | <0.001 |
| 1-2,2-2 | 0.92 | <0.001 |
| P1D,P1S<br>(vertical) | 0.99 | <0.001 |

Well pairs with similar monitoring depths and fairly widely spaced locations were chosen to compute approximately horizontal hydraulic gradients. Similar results were obtained looking at many area gradients from local monitoring wells. Since all of the analyzed wells were strongly correlated and highly covariant, well 3-19 was chosen as a representative example (Table 1).

## Results and discussion

In the one-month data set, there are multiple periods of short-lived but large-magnitude excursions from the mean oxygen concentrations at each of three depths under study within the hyporheic zone (Figure 4). We focus on two 48-hour periods labeled “perturbation A” and “perturbation B”. Dissolved oxygen provides the most sensitive indicator of changing conditions, followed by nitrate concentration. This agrees with prior observations of the typically-oxic nature of both the surface water and shallow groundwater in this region [*Ahmed, et al*., 2012]. Our goal is to understand the observed hydrobiogeochemical dynamics in the context of this hydropeaked river system.

**Figure 4:**
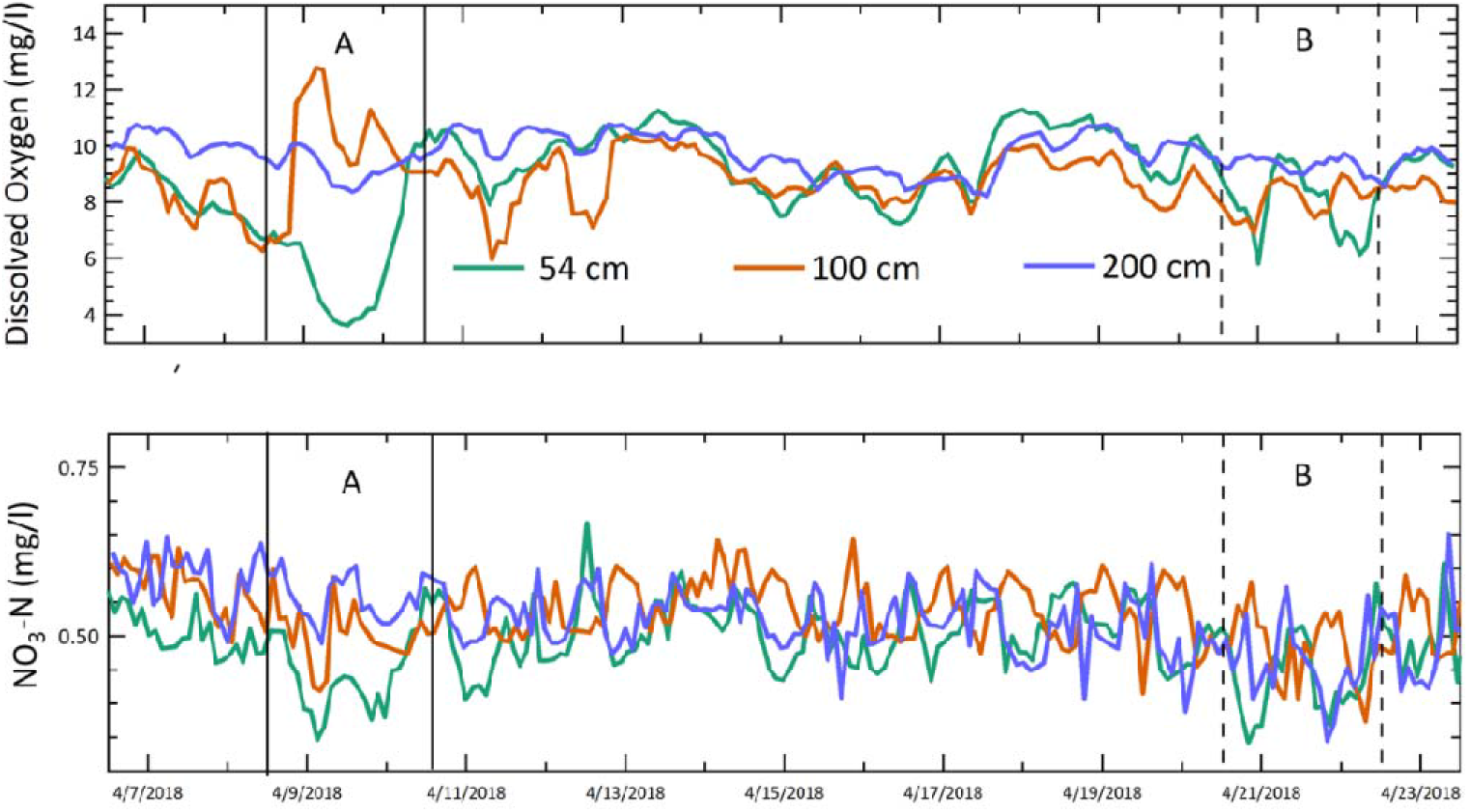
Dissolved oxygen and nitrate (as nitrogen) concentrations at each monitoring depth. The pairs of solid and dashed black vertical lines delineate perturbations A and B.

River stage and hydraulic gradients during the same period are plotted in figure 5. Hydraulic gradient was defined such that positive gradients describe a situation where hydraulic head in the monitoring well was higher than head in the river piezometer, driving water generally toward the river. Negative gradients describe the opposite condition, where water tends to flow away from the river. Note the rapid change in magnitude and direction of the hydraulic gradient during perturbation A in Figure 4. River stage provides the primary control for both the water table elevation in the surficial aquifer as well as horizontal hydraulic gradients. The generally high (though variable) permeability of the surficial aquifer in the area of the study site leads to tight coupling between river and subsurface hydraulics. This allows the river stage to strongly influence potentiometric surface elevations both near the river and farther away at monitoring wells installed approximately 450 meters away from the river. Fluctuating gradients driven by relatively high frequency stage fluctuations are common both in the general area of the field site [*Fritz and Arntzen*, 2007], and also in other river systems [*Watson, et al*., 2018]. These water table elevations in turn control the hydraulic gradient.

**Figure 5:**
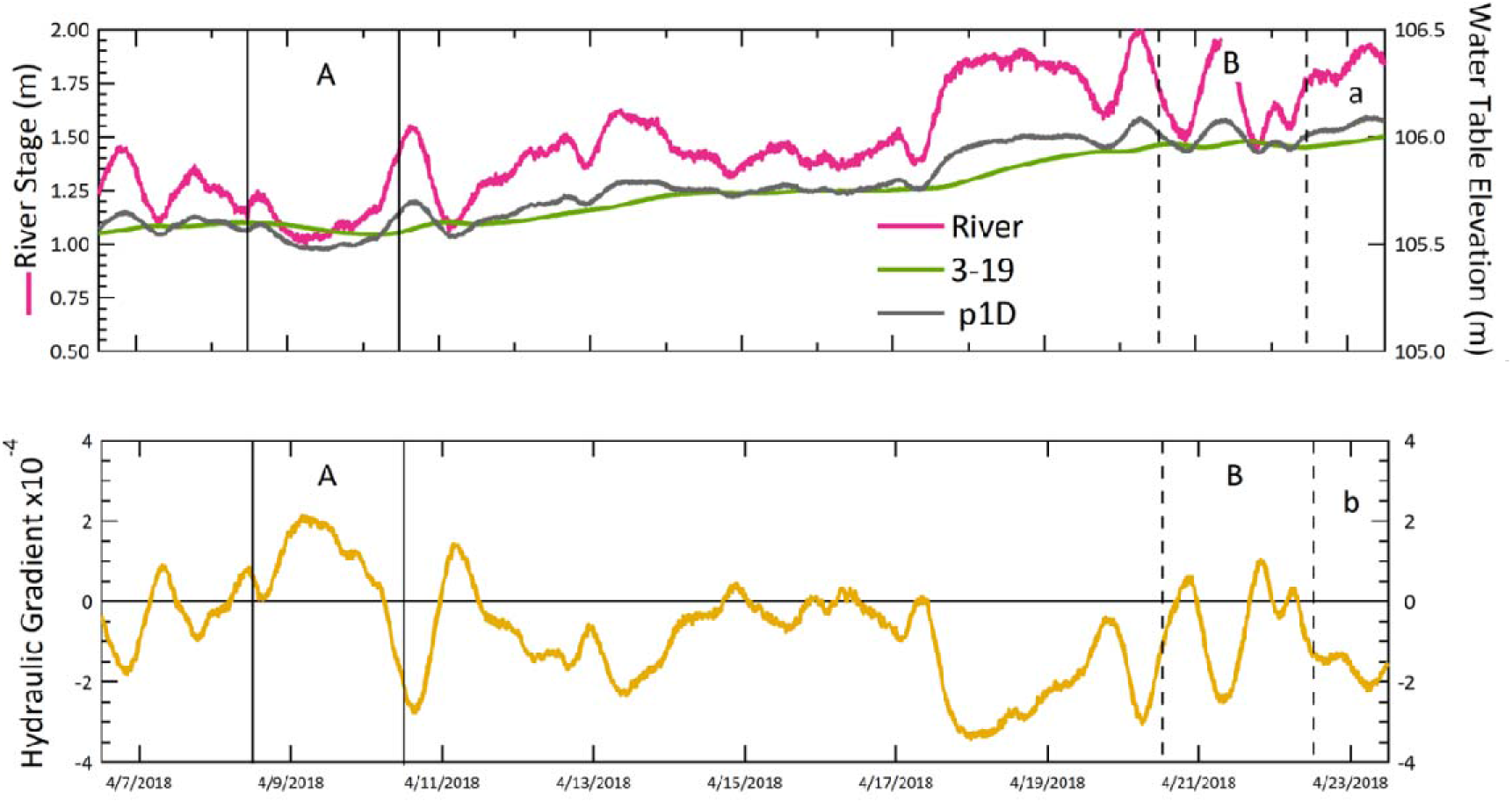
(a) shows the stage height of the river at the monitoring location (referenced to an arbitrary vertical datum), as well as the water table elevations at the two wells chosen to represent the approximately horizontal hydraulic gradient at the study location (referenced to mean sea level). The range of both Y axes is the same. (b) is the hydraulic gradient computed between wells 3-19 and P1D. With a positive gradient, water flows toward the river. The pairs of solid and dashed black vertical lines delineate perturbations A and B.

DO and nitrate varied in response to hydraulic gradient fluctuations (compare the time periods of perturbation A in Figure 4 and Figure 5B respectively). The question arises: does ground and surface water mixing drive the excursions observed during perturbation A, as it is common for variable groundwater/surface water mixing to exert a strong influence on subsurface biogeochemistry [*Fritz and Arntzen*, 2007, *Liu, et al*., 2017, *Stegen, et al*., 2016]? However that does not appear to be the primary process at work at this site. Electrical conductivity consistently agrees with the open channel electrical conductivity of approximately 155 μs/cm (Figure 6). In contrast, the average groundwater conductivity from monitoring well 3-19 was approximately 430 μs/cm and changed very little over the course of the study. While the deepest monitoring location does show slightly higher conductivity than the shallower monitoring depths, this elevation is very small and the overall lack of elevated conductivity at all of the sampling points compared to the river channel implies that little mixing of high-conductivity groundwater with low-conductivity river water is occurring at those locations.

**Figure 6:**
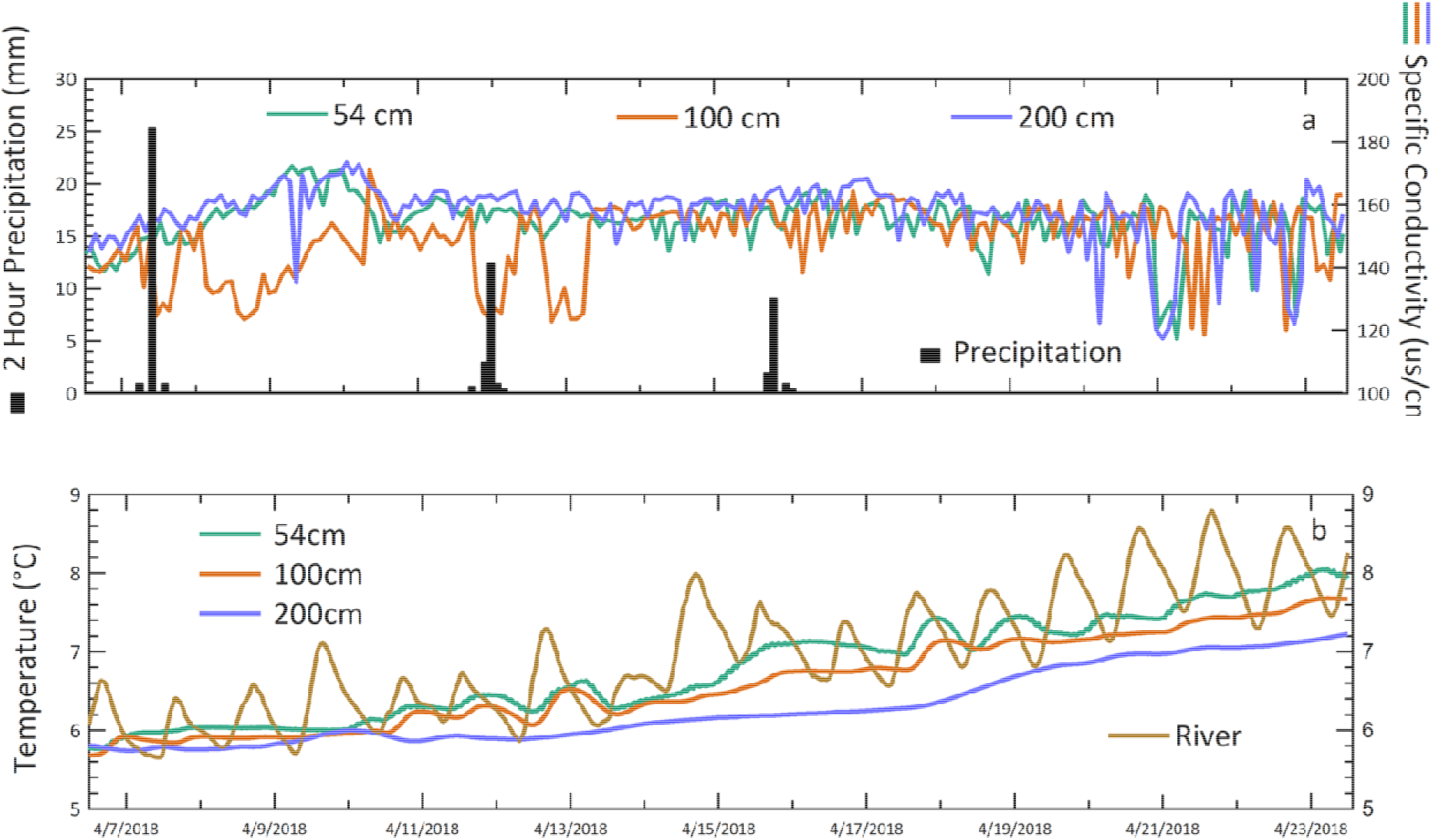
(a) shows the specific conductivity at the three different depths, as well as periods of precipitation. The weather station was located approximately 8 km from the study site, and precipitation patterns in the study area can be extremely localized, so precipitation events should be regarded as approximate. (b) shows the temperature recorded at the three monitoring depths, as well as in the river channel.

Temperature is another driver of biogeochemical transformation rates [*Zheng, et al*., 2016], providing another potential cause of the observed perturbation. However the temperatures recorded at the monitoring points, while correlated with changes in gradient, only varied by about 2°C over the study period, implying that temperature variations within the subsurface, and the corresponding reaction rate variations that would follow, are not the primary drivers of biogeochemical perturbation at this study site.

Having established that both surface/ground water mixing and subsurface temperature are not driving the observed DO perturbations, we analyzed the functional relationship between DO and hydraulic gradient. During our study, DO responds to fluctuations in hydraulic gradient in two distinct forms, linear and non-linear, defining two separate hydrobiogeochemical regimes (Figure 7). In one regime DO is driven by gradient in a broadly linear fashion (Figure 7). This relationship holds true for the bulk of the time series, including perturbation B. The slope of the line varies across the three monitoring depths, but the functional form of the relationship between DO and gradient does not, and it is the functional form of the relationship between DO and hydrologic gradient that defines the hydrobiogeochemical regime. Additionally, the DO concentrations at the three depths tend to vary together in this linear regime, and when they do diverge they diverge in magnitude of perturbation but not direction. This regime is similar to what has been described in a nearby reach of the same river: “Scatterplots of dissolved oxygen … in the hyporheic zone showed a linear trend with river stage” [*Arntzen, et al*., 2006].

**Figure 7:**
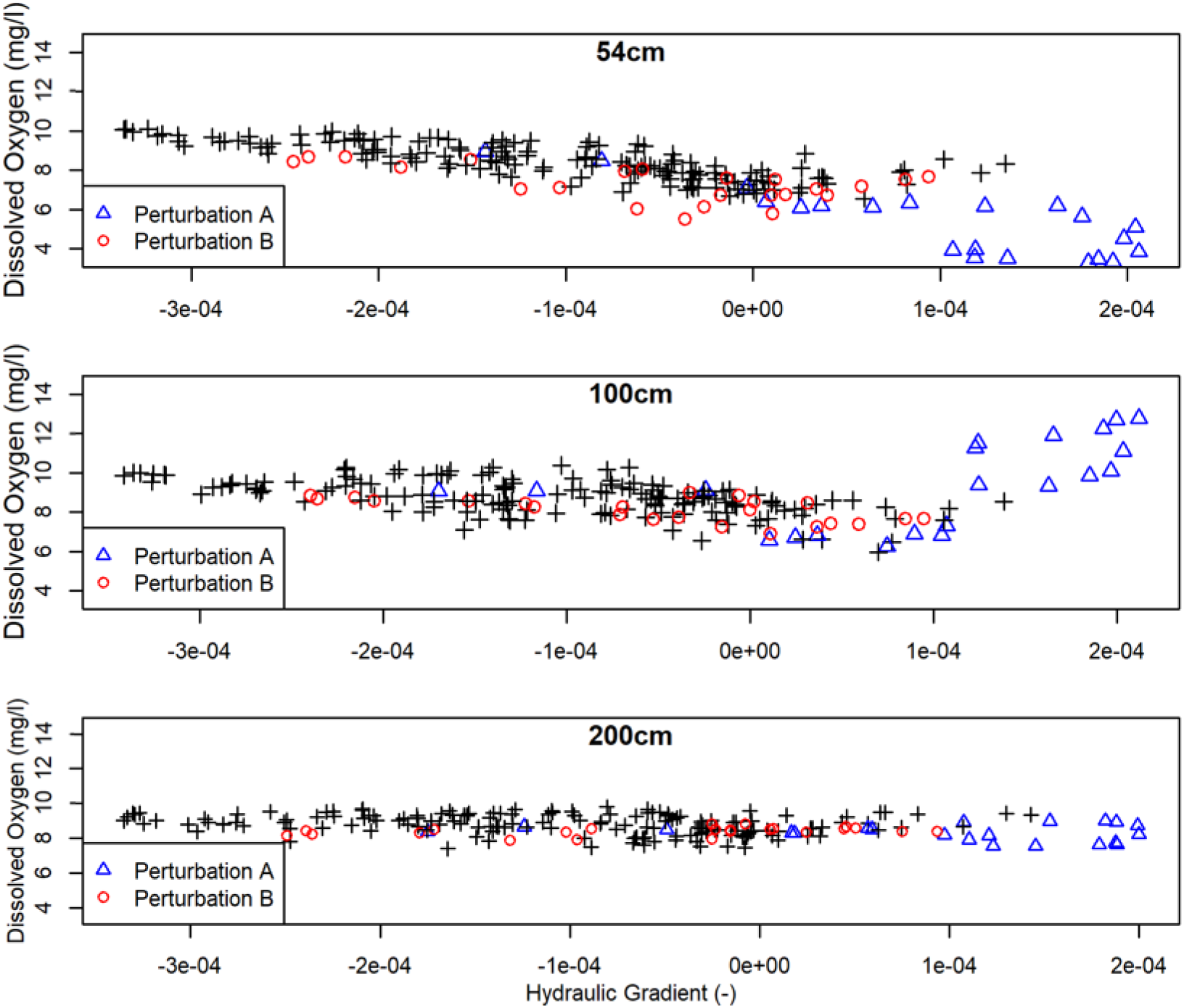
Bivariate plots of dissolved oxygen concentration vs. hydraulic gradient for the entire study time period.

During perturbation A however, we see a non-linear functional form that defines a second hydrobiogeochemical regime. In contrast to the linear regime, we see strong divergence in the DO concentrations at the three depths: the 54cm probe records a strong decrease in DO concentration, while the 100cm probe records an equally strong increase, rising to saturation and staying there for approximately 8 hours (Figure 4), and finally the 200cm probe records little variation. Additionally, we lose the simple perturbation response present in the linear regime and instead observe strong hysteresis at all monitoring depths, despite the variation in magnitude and direction of the apparent DO responses (Figure 8). This hysteretic regime is similar to what *Soulsby, et al*. [2009] observed during a flood event in the river Dee in Scotland.

**Figure 8:**
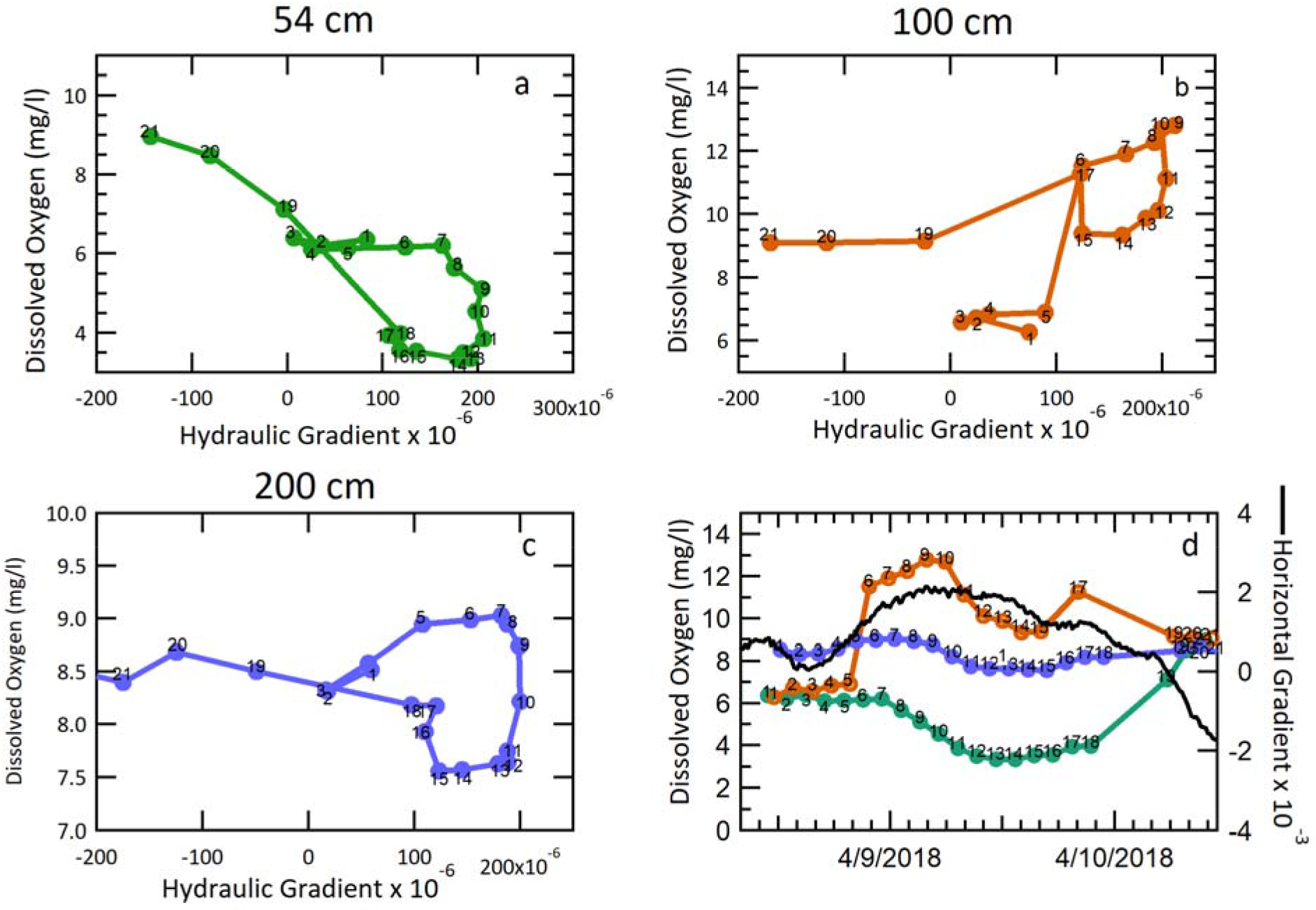
(a-c) are bivariate plots of dissolved oxygen vs. hydraulic gradient for the duration of perturbation A. The datapoints are numbered and connected by a line sequentially in time. The dotted red lines are linear trendlines for the displayed data. The structured loop shape described by these plots indicates a hysteretic process. (d) is the dissolved oxygen time series during perturbation A at all three depths, with the same sequential numbering scheme as (a-c).

Concisely stated, we observe relatively high frequency hydrologic perturbations causing one river to behave like two different rivers with distinct hydrobiogeochemical regimes, only hours apart in time. The time series in Figure 8d shows that between point 5 and 18 the lateral hydraulic gradient switches rapidly from positive to negative, i.e. water flowing towards the river then away from the river. During this event DO displays a hysteretic pattern commencing at point 4, then closing upon itself 28 hours later at point 18. Remarkably, the same hysteric behavior is observed at all three depths.

We hypothesize that the combination of “old” water flushing from the banks and preferential flowpaths leads to DO concentrations that are both dynamic and variable in space. The general hydrologic setting of the field site includes high permeabilities with a strong spatial variability. This, combined with large and rapid stage fluctuations, implies that the mechanism behind the observed change in hydrobiogeochemical regime may be residence time. Perturbation A takes place during a local minimum river stage (Figure 5), which generates a period of relatively strong positive (toward-river) gradient that lasts for 40 hours. Such a period of toward-river gradient may allow river water that has been trapped deep in the banks for some time to be transported from the banks into the riverbed, passing by the monitoring locations. This water would have had longer residence time in the bank, allowing depletion in dissolved oxygen. However, the situation is clearly not that simple, as the DO variation across the monitoring depths differs in both magnitude and direction. As the subsurface here contains significant spatial heterogeneity, preferential flowpaths are sure to exist. The conductivity data alludes to this situation as well, with the conductivity at the 100cm monitoring point dropping significantly immediately after several rain events (Figure 6), while similar declines at the 54cm and 200cm monitoring points were not observed. Additionally, the gradient correlates worse with specific conductivity at 100cm than at the other two depths (Table S2). Both pieces of evidence imply that the low-conductivity rain water has preferential access to the middle monitoring point. Implicit in this proposed conceptual model of the system is predominantly horizontal subsurface flow.

## Conclusions and implications

Our study shows a clear example of short-term stage fluctuations driving the system to abruptly shift between linear and hysteretic hydrobiogeochemical regimes across short time scales in the location sampled. A combination of hydrologic conditions at our study site causes the system to ‘tip’ from one hydrobiogeocheimcal regime to another. This threshold-like behavior makes the overall hydrobiogeochemical system difficult to predict, even with a relatively high frequency data. It is unlikely that our setting is unique, and thus the potential exists for such threshold behavior in hydrobiogeochemical regime in any system experiencing short-term stage dynamics. While the stage fluctuation in our system is driven by a hydroelectric dam, short-term stage dynamics are present in many other types of river corridor systems, including those influenced by strong evapotranspiration, tidal forces, municipal or agricultural withdrawals, or other similar processes.

The subsurface component of river corridor systems can be important to wider scale fluxes of nutrients and organic material [*Krause, et al*., 2011, *Wondzell*, 2011]. Short-term stage dynamics are common across river corridors, and to the extent that our results can be generalized, they imply that the potential for threshold behavior presents a significant challenge to the estimation of cumulative influences of subsurface processes from reach to earth system scales.

While multi-parameter spatial and temporal monitoring in the subsurface is challenging, it is required to observe impacts of disturbances on hydrobiogeochemical regimes. The biogeochemical observables in this system show significant changes occurring on the time scale of hours. Importantly, constraining a numeric or even conceptual model of such a complicated and dynamic system relies on a diverse set of measurements, including DO, conductivity, temperature, hydraulic gradient and many more. Some species, such as DO, serve as key indicators of a particular regime, whereas measurements that show little dynamic character or poor correlations (such as ORP) are nonetheless important in constraining modelling efforts. The variable permeability of the study site study suggests that data from a greater spatial extent will aid in further developing our understanding of the different HBGC regimes. The identification and study of hydrobiogeochemical regimes and inter-regime dynamics thus requires instrumentation that can gather a diverse set of measurements from a large number of sampling sites on rapid time scales. The stabilization time of the present system (about 40 minutes) implies that increasing the spatial resolution of the system will likely require changes to the mechanical function of the system, parallel installations of the pumping and analysis systems, or potentially an increased deployment of in-situ measurement devices. This difficulty can be at least partly ameliorated through by using spatially distributed realtime or rapid-response measurements, such as electrical resistance tomography, to target a specific subset of useful monitoring locations for sampling at any given time.

## Acknowledgments

This research is supported by the U.S. Department of Energy (DOE), Office of Biological and Environmental Research (BER), as part of BER’s Subsurface Biogeochemical Research Program, (SBR). This contribution originates from the SBR Scientific Focus Area (SFA at the Pacific Northwest National Laboratory (PNNL). This research is supported under SBR award DE-SC0018042. A portion of the research described in this paper was conducted under the Laboratory Directed Research and Development Program at Pacific Northwest National Laboratory, a multiprogram national laboratory operated by Battelle for the U.S. Department of Energy. The authors are grateful for the support of the Linus Pauling Distinguished Postdoctoral Fellowship program. This material is based upon work supported by the US Dept. of Energy, Office of Science, Small Business Initiative Research, under award number DE-SC0017129. The dataset for this publication is hosted publicly at the following location: – URL to be inserted--

